# Binding Ligands that Straddle an Important Contact Site on the RBD of the Covid-19 Spike Protein

**DOI:** 10.1101/2020.08.03.234989

**Authors:** Abraham Boyarsky

## Abstract

The receptor binding domain (RBD) of the spike protein of the Covid-19 virus is responsible for attachment to human ACE2. A number of recent articles have studied monoclonal antibody blocking [8-11] and peptide inhibitors [12-16] of the Covid-19 virus. Here we report virtual ligand-based screening that targets pockets on each side of an important binding site with residues 502-504 on the RBD that contact residues 353-357 [15] of hACE2. These ligands are intended as pre-exposure therapy for Covid-19 infection.

## Introduction

Since December 2019, the Covid-19 outbreak has caused more than 10 million infections and more than a half million deaths. The RBD of the spike protein of Covid-19, a sequence of 194 residues, interacts directly and tightly with hACE2. To access host cells, the Covid-19 virus relies on its spike protein, which contains a receptor binding segment S_1_ and a fusion segment S_2_ [6]. The S_1_ segment contains the receptor binding domain (194 amino acids) that binds to the host cell receptor. The human ACE2 is the critical receptor for entry into cells [7,8].

Although there are a number of Covid-19 strains circulating globally (2), the RBD sequence seems to be mostly conserved. A number of recent articles have studied monoclonal antibody blocking [8-11] and peptide inhibitors [12-16] of the Covid-19 virus. In this note we study the surface of RBD for pockets and screen for ligands that target these sites as potential drugs against the Covid-19 virus.

The PBD file 6YLA contains the crystal structure of the SARS-CoV-2 spike RBD bound with hACE2. Figure 1 displays the interface of RBD and hACE2. A segment of an alpha helix is at the forefront of the interface with residues 502-504, much as a scaffold leading the advance on hACE2 (Figure 1). These residues were also identified in [13] as a significant anchoring segment. The aim of this study is to search for small molecule inhibitors that have strong affinity to pockets on either side of the residues of 502-504 of RBD and could interfere with the interaction of the RBD-ligand complex with hACE2. In this note we provide hits that are known drugs that may be repurposed as prophylactic and therapeutic interventions against Covid-19 infection.

**Figure 1.**
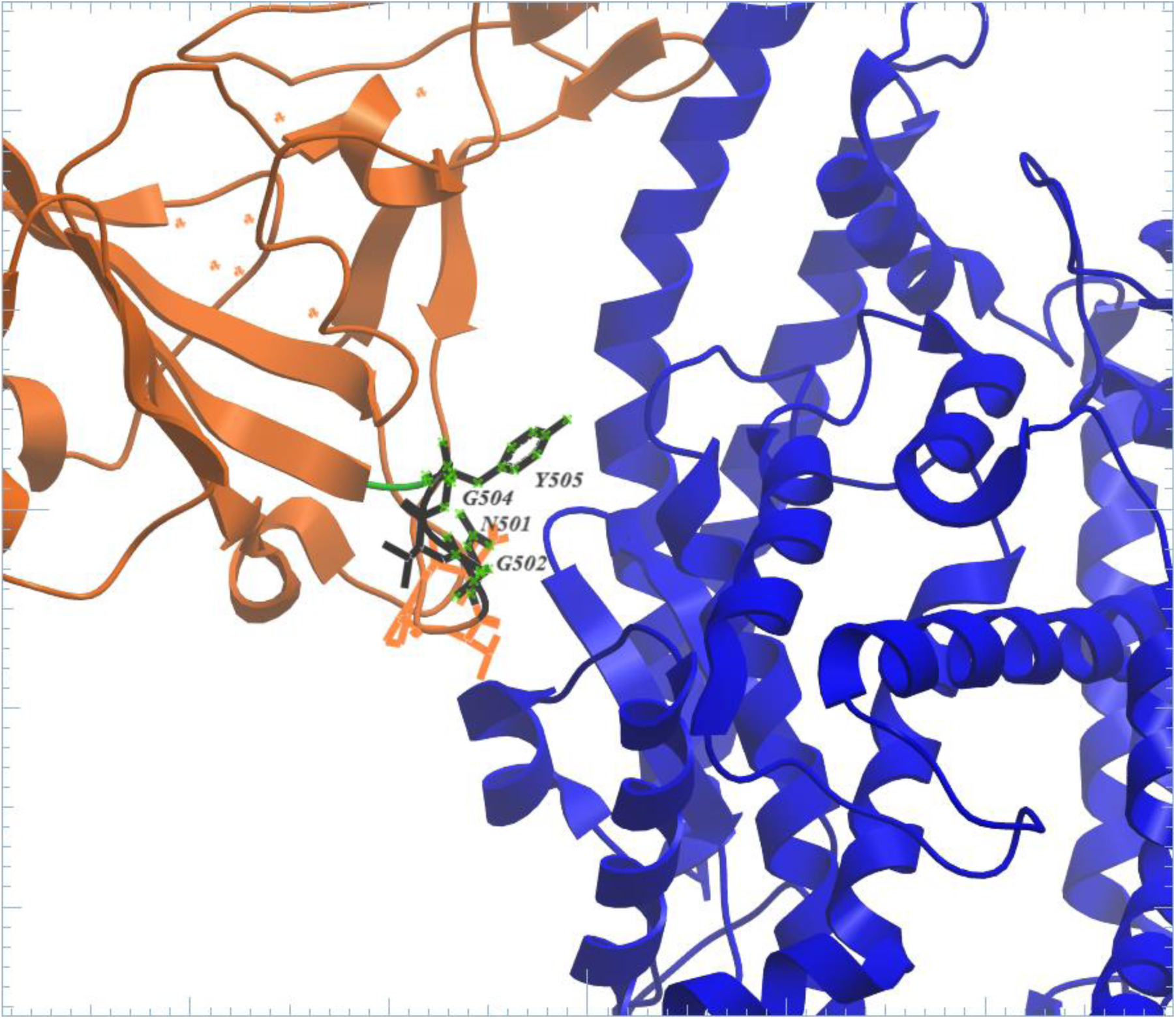
Interaction interface of RBD (brown)-hACE2 (blue). Residues on advancing segment labelled.

We are targeting a protein-protein interaction site and since protein-protein interfaces are generally flat, there are usually few good cavities for small molecules. The two pockets in Figure 2 surround the binding site G502 V503 G504. The pocket includes RBD residues R403, Y449, Y453, Q493, S494, Y495, G496, F497, Q498, N501 and Y505. It is expected that chemical binding to pockets would affect the structure of the binding interface and interfere with the binding of RBD-ligand complex to hACE2.

**Figure 2.**
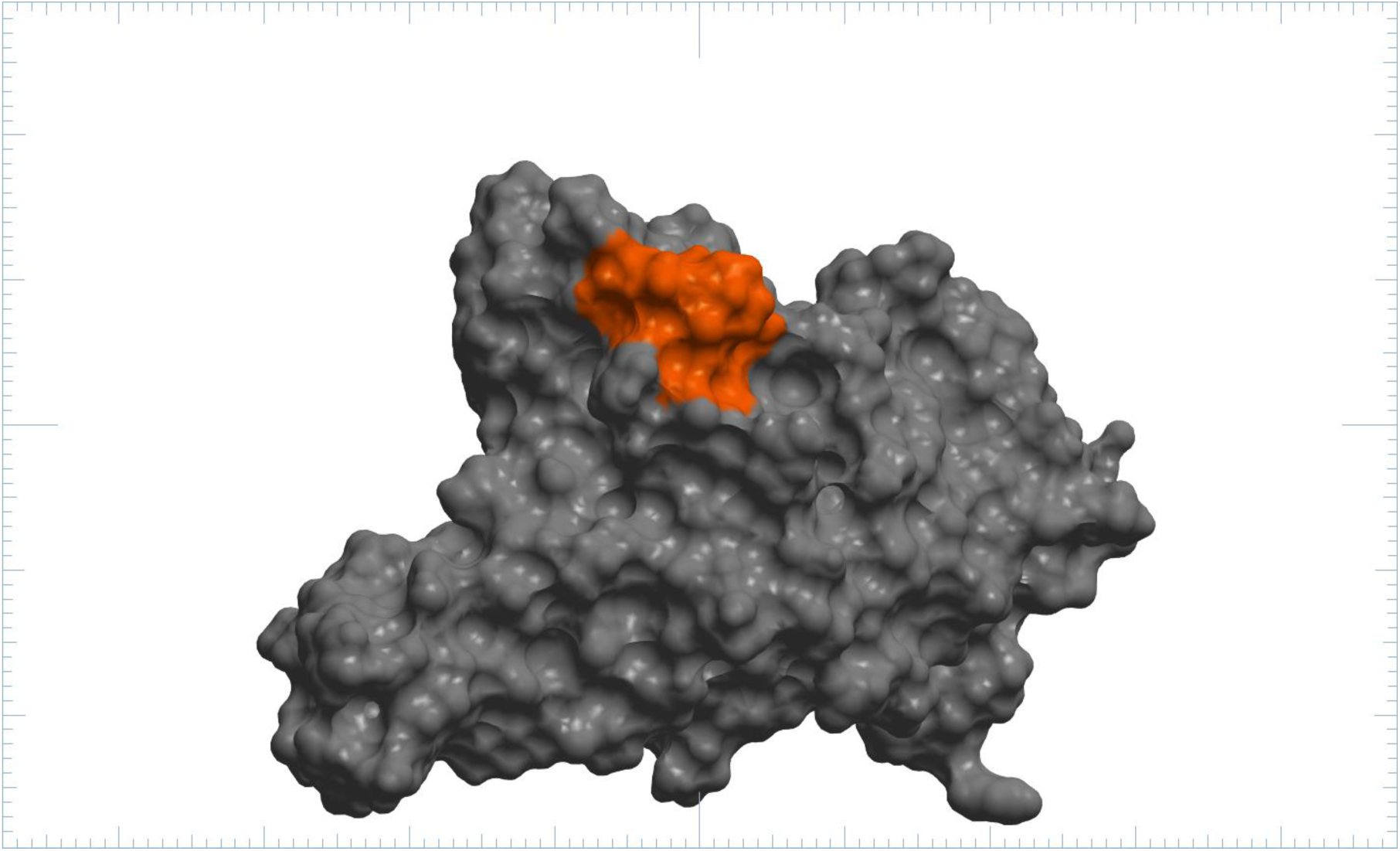
Surface of RBD (grey). Residues G502-V503-G504 (orange).

## VLS Methods

For virtual ligand screening (VLS) we used the crystal structure of the Covid-19 receptor binding domain (RBD) with PDB code 6YLA. All molecules were removed except for the “e” chain of the structure. Using the crystal structure of the RBD-ACE2 complex, PDB 6LZG [1], the region G502-V503-G504 of the RBD was selected as the target surrounded by the RBD residues R403, Y449, Y453, Q493, S494, Y495, G496, F497, Q498, N501 and Y505.

The MolCart Compounds 2020 database (http://www.molsoft.com/screening.html) which contains ∼10.2 M commercially available chemicals was then set up to screen against the pocket using MolSoft’s ICM-VLS docking method [2,3].

For the virtual screen five types of interaction potentials represented the binding pocket: (i) van der Waals potential for a hydrogen atom probe; (ii) van der Waals potential for a heavy-atom probe (generic carbon of 1.7 Å radius; (iii) optimized electrostatic term; (iv) hydrophobic terms; and (v) loan-pair-based potential, which reflects directional preferences in hydrogen bonding. The energy terms are based on the all-atom vacuum force field ECEPP/3 with appended terms to account for solvation free energy and entropic contribution. Conformational sampling is based on the biased probability Monte Carlo (BPMC) procedure [4], which randomly selects a conformation in the internal coordinate space and then makes a step to a new random position independent of the previous one but according to a predefined continuous probability distribution. It has also been shown that after each random step, full local minimization greatly improves the efficiency of the procedure. The ICM program relies on global optimization of the entire flexible ligand in the receptor field and combines large-scale random moves of several types with gradient local minimization and a search history mechanism.

The scoring function gives a good approximation of the binding free energy between a ligand and a receptor and is a function of different energy terms based on a force-field. The ICM scoring function [5] is weighted according to the following parameters (i) internal force-field energy of the ligand, (ii) entropy loss of the ligand between bound and unbound states, (iii) ligand-receptor hydrogen bond interactions, (iv) polar and non-polar solvation energy differences between bound and unbound states, (v) electrostatic energy, (vi) hydrophobic energy, and (vii) hydrogen bond donor or acceptor desolvation. The lower the ICM score, the higher the chance chance the ligand is a binder.

Atomic Property Field (APF) is a 3D pharmacophoric potential implemented on a grid and was developed by MolSoft [6]; it is used to select a diverse number of chemicals for testing. APF can be generated from one or multiple ligands, and seven components are assigned from empiric physico-chemical properties (hydrogen bond donors, acceptors, Sp2 hybridization, lipophilicity, size, electropositive/negative charge). In the case of docking hit clustering, the APF field is used to compare 3D distributions of these properties for pairs of ligands in their docked poses within the pocket. This allows to cluster together compounds that dock similarly with similar properties.

The APF clustering of the VLS hitlist helps choose a diverse set of compounds for testing or helps easily locate a series of interesting compounds. The tree and table are fully interactive therefore clicking on a compound in the tree will highlight the compound in the table and display the complex.

## VLS Results

After filtering out chemicals with a high ToxScore and predicted to be PAINS (see [7]) the final hitlist has 10159 chemicals. The chemicals in the hitlist are ranked by SCORE, when choosing chemicals for testing we recommend selecting based on Score (the lower the better). The clustering helps in the selection of a diverse set. The more compounds that are tested the higher the chances of success. The chemical vendors are listed in the column called “vendor” along with the chemical ID number.

## Data collection with SPR

The goal is to determine binding affinity KD and kinetic parameters ka, kd of Recombinant COVID-19 S(RBD) protein with 6 compounds.

## Methods and Materials

### Samples

- Recombinant COVID-19 S(RBD) protein (Cat# nCoVS-125V) is provided by Profacgen.
- C34H30N4O5, C18H24N6O3 and C16H22N6O2 are purchased from Chembridge.
- C28H20N2O7 is purchased from Chemdiv.
- C24H20N4O5 and C16H13IN4O2S are purchased from Enamine.

### Reagents and Consumables

- Reagents and use of instrument were provided by Profacgen.

### Equipment

- PlexAray HT (Plexera Bioscience, Seattle, WA, US).

## Methods

The bare gold-coated (thickness 47 nm) PlexArray Nanocapture Sensor Chip (Plexera Bioscience, Seattle, WA, US) was prewashed with 10× PBST for 10 min, 1× PBST for 10 min, and deionized water twice for 10 min before being dried under a stream of nitrogen prior to use. Various concentrations of biotinylated recombinant COVID-19 S(RBD) protein dissolved in water were manually printed onto the Chip with Biodo bioprinting at 40% humidity via biotin-avidin conjugation. Each concentration was printed in replicate, and each spot contained 0.2 µL of sample solution. The chip was incubated in 80% humidity at 4°C for overnight, and rinsed with 10× PBST for 10 min, 1× PBST for 10 min, and deionized water twice for 10 min. The chip was then blocked with 5% (w/v) non-fat milk in water overnight, and washed with 10× PBST for 10 min, 1× PBST for 10 min, and deionized water twice for 10 min before being dried under a stream of nitrogen prior to use. SPRi measurements were performed with PlexAray HT (Plexera Bioscience, Seattle, WA, US). Collimated light (660 nm) passes through the coupling prism, reflects off the SPR-active gold surface, and is received by the CCD camera. Buffers and samples were injected by a non-pulsatile piston pump into the 30 µL flowcell that was mounted on the coupling prim. Each measurement cycle contained four steps: washing with PBST running buffer at a constant rate of 2 µL/s to obtain a stable baseline, sample injection at 5 µL/s for binding, surface washing with PBST at 2 µL/s for 300 s, and regeneration with 0.5% (v/v) H3PO4 at 2 µL/s for 300 s. All the measurements were performed at 25°C. The signal changes after binding and washing (in AU) are recorded as the assay value.

### Kinetics fitting and analysis

Selected protein-grafted regions in the SPR images were analyzed, and the average reflectivity variations of the chosen areas were plotted as a function of time. Real-time binding signals were recorded and analyzed by Data Analysis Module (DAM, Plexera Bioscience, Seattle, WA, US). Kinetic analysis was performed using BIA evaluation 4.1 software (Biacore, Inc.).

### Experimental results

The experimental results are collected in Table 1 below. The binding affinity graphs are shown in Figures 5-10.

**Table 1.** The binding affinity KD and kinetic parameters ka, kd of Recombinant COVID-19 S(RBD) protein with 6 compounds.

| Compound | KD | k<sub>a</sub> | k<sub>d</sub> |
| --- | --- | --- | --- |
| C24H20N4O5 | $4.14 \times 10^{-5}$ M | $1.58 \times 10^2 \text{ M}^{-1} \cdot \text{s}^{-1}$ | $6.54 \times 10^{-3} \text{ s}^{-1}$ |
| C16H13IN4O2S | $1.78 \times 10^{-4}$ M | $0.85 \times 10^2 \text{ M}^{-1} \cdot \text{s}^{-1}$ | $1.51 \times 10^{-2} \text{ s}^{-1}$ |
| C28H20N2O7 | $1.02 \times 10^{-6}$ M | $3.27 \times 10^2 \text{ M}^{-1} \cdot \text{s}^{-1}$ | $3.34 \times 10^{-4} \text{ s}^{-1}$ |
| C34H30N4O5 | $3.01 \times 10^{-5}$ M | $6.55 \times 10^2 \text{ M}^{-1} \cdot \text{s}^{-1}$ | $1.97 \times 10^{-2} \text{ s}^{-1}$ |
| C18H24N6O3 | $8.36 \times 10^{-7}$ M | $7.12 \times 10^2 \text{ M}^{-1} \cdot \text{s}^{-1}$ | $5.95 \times 10^{-4} \text{ s}^{-1}$ |
| C16H22N6O2 | $1.67 \times 10^{-5}$ M | $0.97 \times 10^2 \text{ M}^{-1} \cdot \text{s}^{-1}$ | $1.62 \times 10^{-3} \text{ s}^{-1}$ |

**Figure 3.**
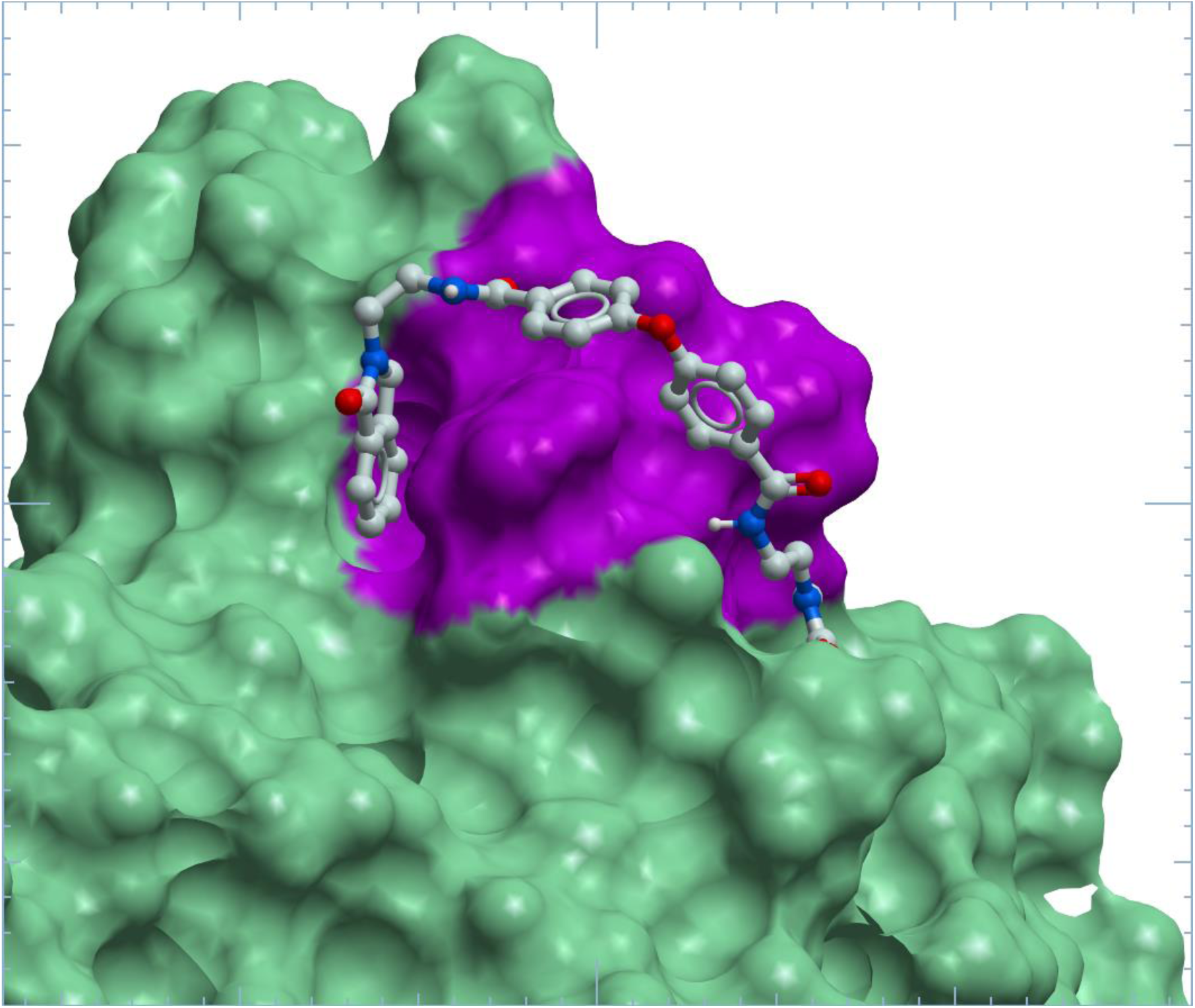
Stick and ball presentation of ligand C34H30N4O5 straddling residues G502-V503-G504 (purple).

**Figure 4.**
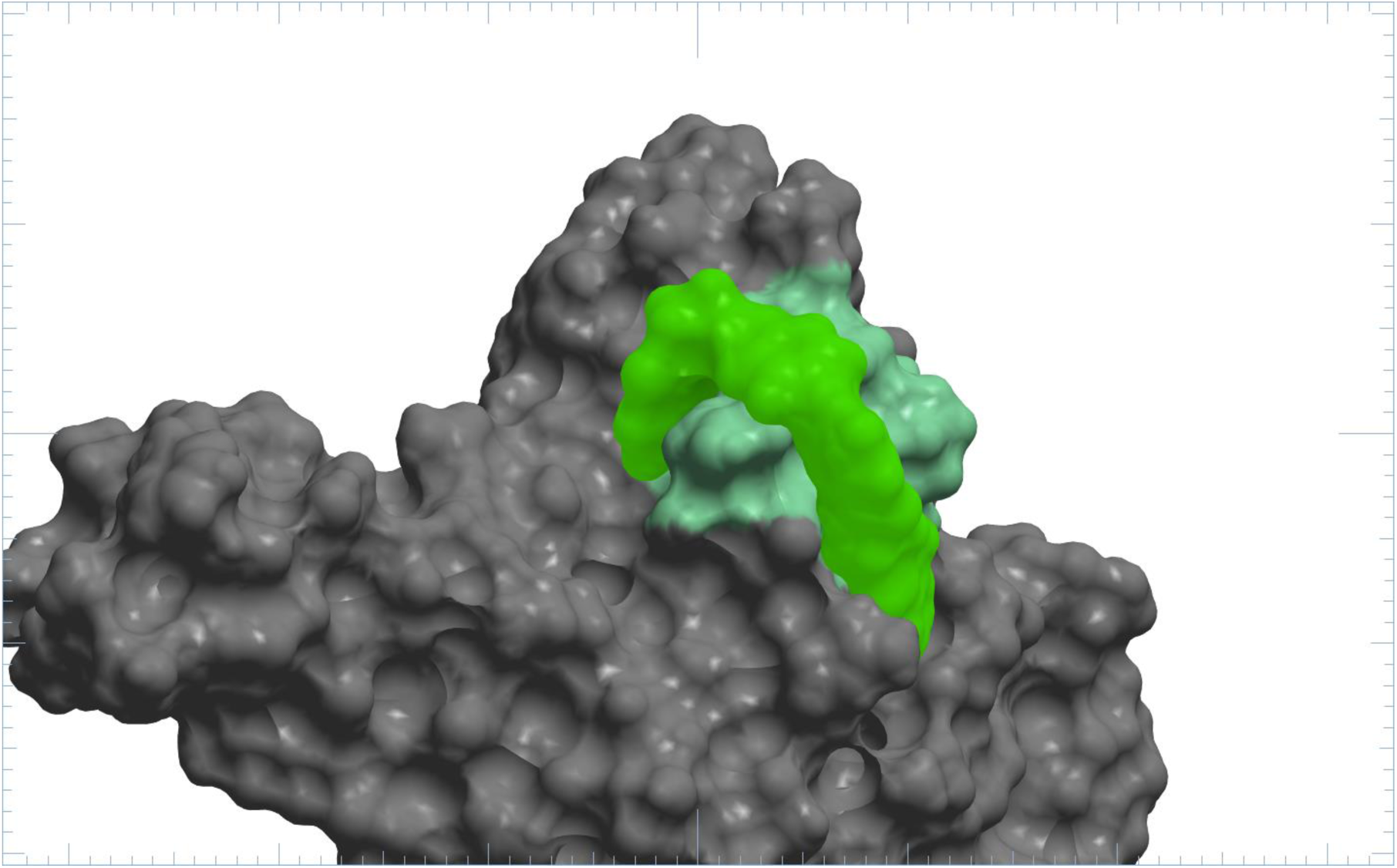
Ligand C24H20N4O5 (green) straddling G502-V503-G504 (aquarine) and bonded to RBD (grey).

**Figure 5.**
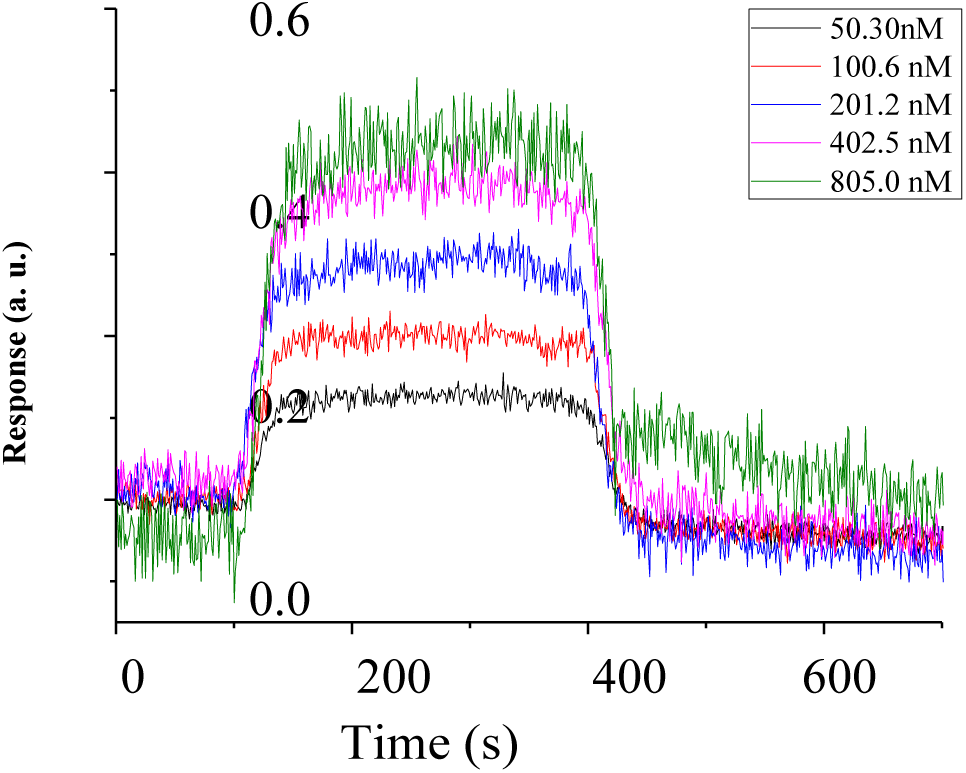
Kinetics fitting curve of Recombinant COVID-19 S(RBD) protein to C24H20N4O5 compound.

## Discussion

We study the surface of RBD for ligand-binding pockets. We focussed on the important residues 502-504 which are on an alpha helix segment at the forefront of the interface of RBD with hACE2. On either side of these residues pockets were identified at R403, Y449, Y453, Q493, S494, Y495, G496, F497, Q498, N501, Y505. Then 10.2 million ligands were screened and their binding affinities scored using Molsoft software. A number of ligands fit into these pockets and straddled the residues 502-504, acting as a guard rail and potentially blocking these and other neighboring residues from binding with hACE2.

Laboratory work was performed to test the binding affinity of the selected ligands to RBD. The binding affinity curves are biphasic, and imply one side is a stronger fit than the other. C28H20N2O7 binds to two sites on the spike protein. One is very tight and the second is weaker. C18H24N6O3 and C16H22N6O2 also bind to two sites on the spike protein. One is very tight and the second is weaker.

## Acknowledgements

The author thanks Dr. Andrew Orr of Molsoft for his computational assistance and to his wife Eva for editorial assistance.

---

**Table 2:**
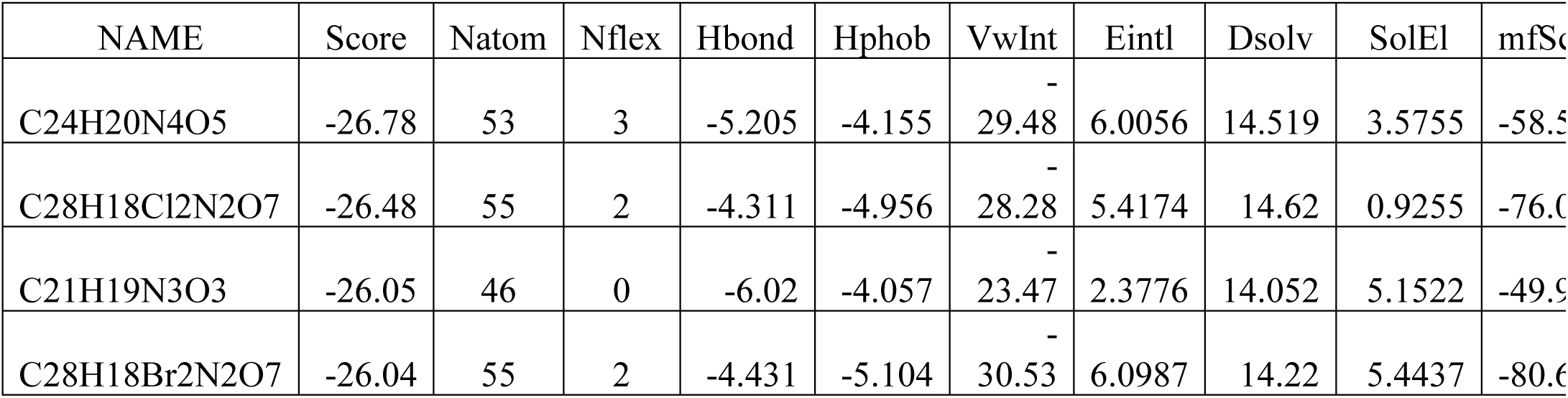
Top hits of ligands that span G502-V503-G504 of RBD.

Columns in Table 1:

Name is the chemical formula.

Score is the VLS score – the lower the score the better the predicted binding.

Natom is the number of atoms in docked ligand.

Nflex is the number of rotatable torsions.

Hbond is Hydrogen Bond energy.

Hphob is the hydrophobic energy in exposing a surface to water.

VwInt is the van der Waals interaction energy (sum of gc and gh van der Waals).

Eintl is internal conformation energy of the ligand.

Dsolv is the desolvation of exposed h-bond donors and acceptors.

SolEl is the solvation electrostatic energy change upon binding.

mfScore is the potential of mean force score.

## Kinetics Fitting Curves

Evaluation of the binding affinity and affinity parameters of cecombinant COVID-19 S(RBD) protein to C24H20N4O5 compound.

The equilibrium dissociation constant (KD Value) was 4.14×10^−5^ M. (ka= 1.58×10^2^ M^-^ 1·s^-1^, kd=6.54×10^−3^s^-1^).

Evaluation of the binding affinity and affinity parameters of Recombinant COVID-19 S(RBD) protein to C16H13IN4O2S compound.

**Figure 6.**
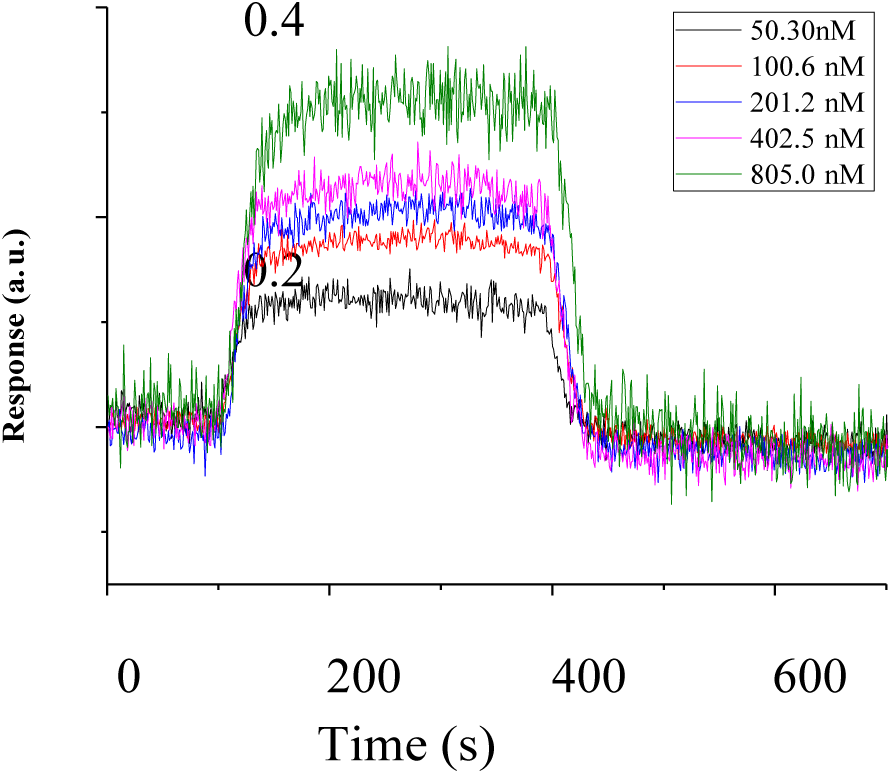
Kinetics fitting curve of Recombinant COVID-19 S(RBD) protein to C16H13IN4O2S compound.

The equilibrium dissociation constant (KD Value) was 1.78×10^−4^ M. (ka= 0.85×10^2^ M^-^ 1·s^-1^, kd=1.51×10^−2^s^-1^).

Evaluation of the binding affinity and affinity parameters of Recombinant COVID-19 S(RBD) protein to C28H20N2O7 compound.

**Figure 7.**
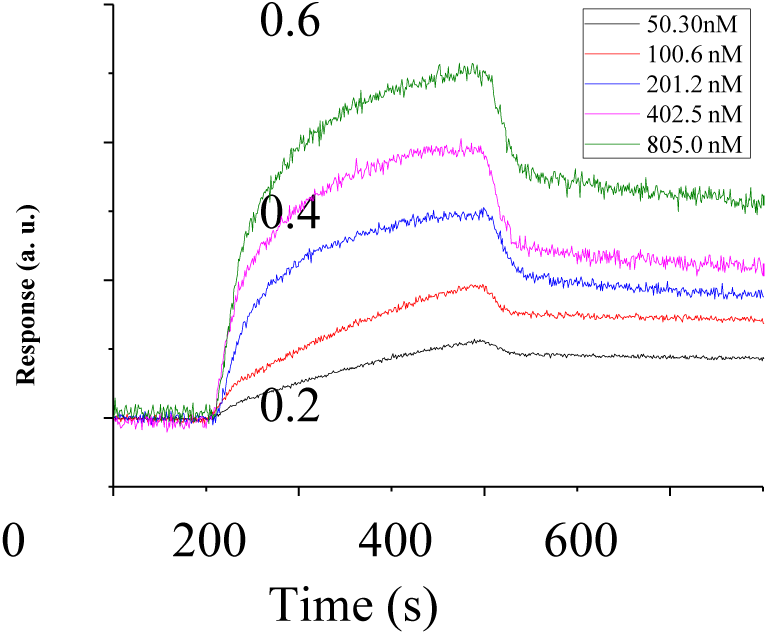
Kinetics fitting curve of Recombinant COVID-19 S(RBD) protein to C28H20N2O7 compound.

The equilibrium dissociation constant (KD Value) was 1.02×10^−6^ M. (ka= 3.27×10^2^ M^-^ 1·s^-1^, kd=3.34×10^−4^s^-1^).

Evaluation of the binding affinity and affinity parameters of Recombinant COVID-19 S(RBD) protein to the C34H30N4O5 compound.

**Figure 8.**
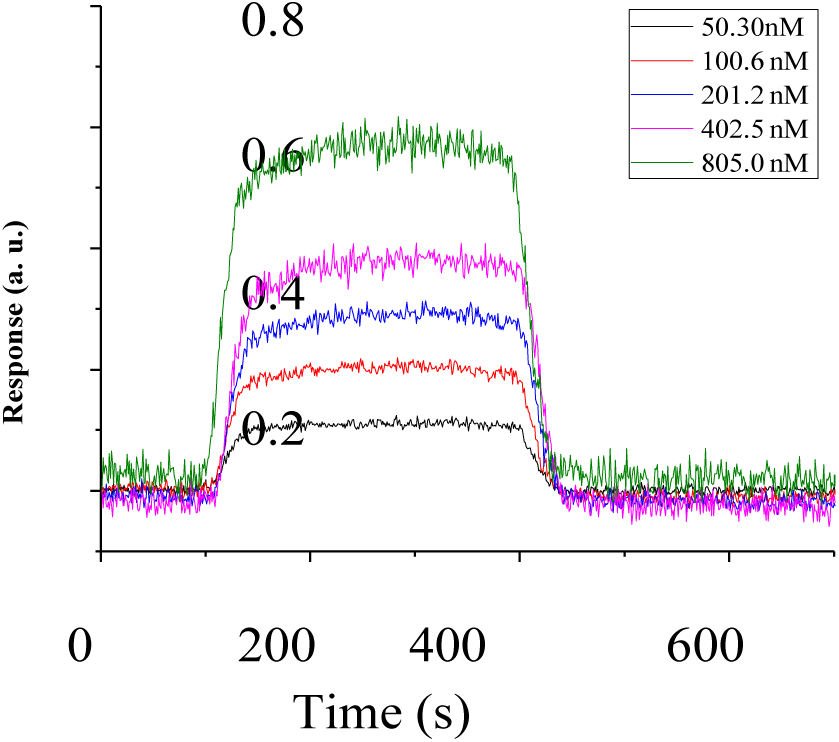
Kinetics fitting curve of Recombinant COVID-19 S(RBD) protein to C34H30N4O5 compound.

The equilibrium dissociation constant (KD Value) was 3.01×10^−5^ M. (ka= 6.55×10^2^ M^-^ 1·s^-1^, kd=1.97×10^−2^s^-1^).

Evaluation of the binding affinity and affinity parameters of Recombinant COVID-19 S(RBD) protein to C18H24N6O3 compound.

**Figure 9.**
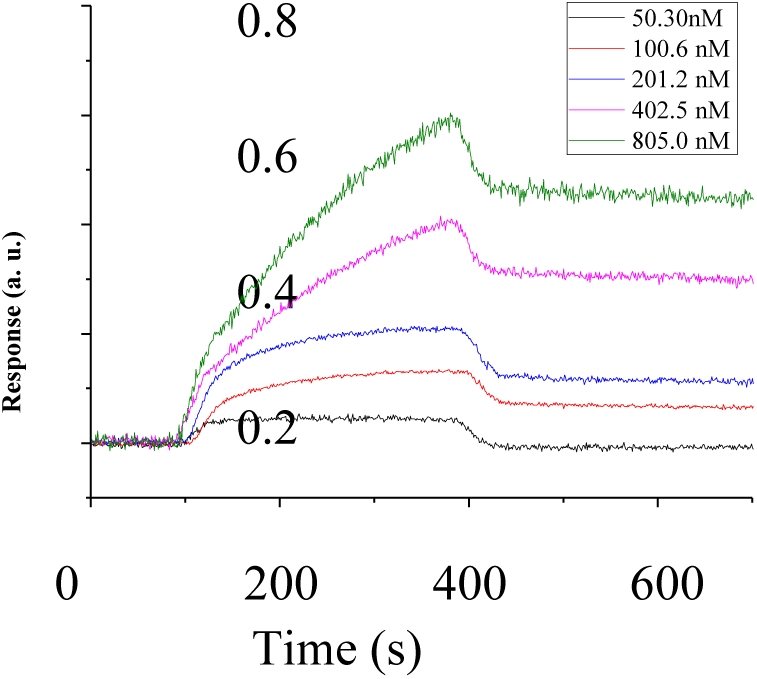
Kinetics fitting curve of Recombinant COVID-19 S(RBD) protein to C18H24N6O3 compound.

The equilibrium dissociation constant (KD Value) was 8.36×10^−7^ M. (ka= 7.12×10^2^ M^-^ 1·s^-1^, kd=5.95×10^−4^s^-1^).

Evaluation of the binding affinity and affinity parameters of Recombinant COVID-19 S(RBD) protein to C16H22N6O2 compound.

**Figure 10.**
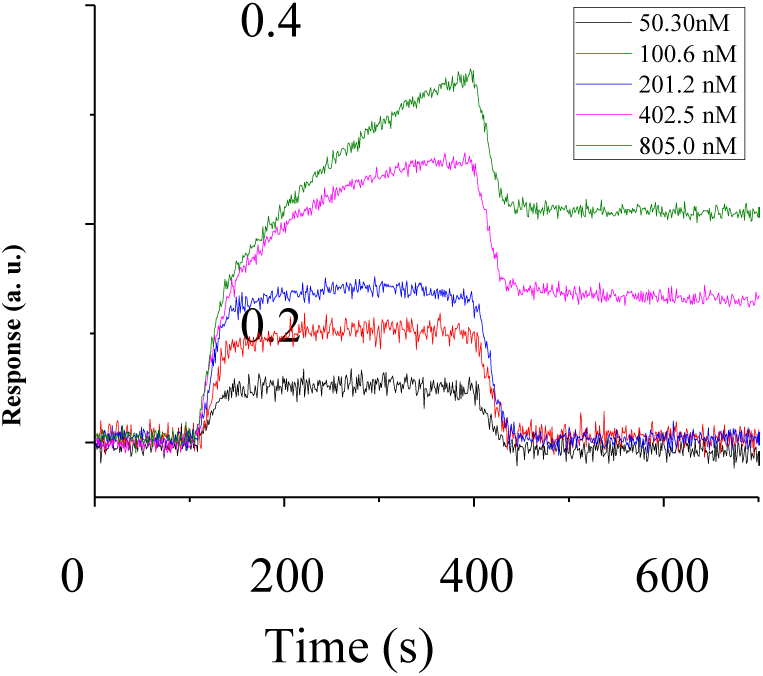
Kinetics fitting curve of Recombinant COVID-19 S(RBD) protein to C16H22N6O2 compound.

The equilibrium dissociation constant (KD Value) was 1.67×10^−5^ M. (ka= 0.97×10^2^ M^-^·s^-1^, kd=1.62×10^−3^s^-1^).

